# A synthetic gene circuit for measuring autoregulatory feedback control

**DOI:** 10.1101/029116

**Authors:** Miquel Àngel Schikora Tamarit, Carlos Toscano-Ochoa, Júlia Domingo Espinós, Lorena Espinar, Lucas B. Carey

## Abstract

Auto regulatory feedback loops occur in the regulation of molecules ranging from ATP to MAP kinases to zinc. Negative feedback loops can increase a system’s robustness, while positive feedback loops can mediate transitions between cell states. Recent genome-wide experimental and computational studies predict hundreds of novel feedback loops. However, not all physical interactions are regulatory, and many experimental methods cannot detect self-interactions. Our understanding of regulatory feedback loops is therefore hampered by the lack of high-throughput methods to experimentally quantify the presence, strength, and temporal dynamics of auto regulatory feedback loops. Here we present a mathematical and experimental framework for high-throughput quantification of feedback regulation, and apply it to RNA binding proteins (RBPs) in yeast. Our method is able to determine the existence of both direct and indirect positive and negative feedback loops, and to quantify the strength of these loops. We experimentally validate our model using two RBPs which lack native feedback loops, and by the introduction of synthetic feedback loops. We find that the the RBP Puf3 does not natively participate in any direct or indirect feedback regulation, but that replacing the native 3’UTR with that of COX17 generates an auto-regulatory negative feedback loop which reduces gene expression noise. Likewise, the RBP Pub1 does not natively participate in any feedback loops, but a synthetic positive feedback loop involving Pub1 results in increased expression noise. Our results demonstrate a synthetic experimental system for quantifying the existence and strength of feedback loops using a combination of high-throughput experiments and mathematical modeling. This system will be of great use in measuring auto-regulatory feedback by RNA binding proteins, a regulatory motif that is difficult to quantify using existing high-throughput methods.

## INTRODUCTION

Homeostatic maintenance of cell state and transitions between states are often mediated by sets of feedback loops^1^. These loops can be positive or negative, and either directly auto-regulatory or indirect, acting through any number of intermediate genes. Both positive and negative feedback loops are used by both organisms and synthetic biologists to perform a wide range of tasks, e.g., ATP biosynthesis^2^, MAPK signaling^3^ and zinc homeostasis^4^. Genome-wide experimental measurements and computational predictions of protein-protein and protein-RNA interactions suggest the existence of thousands of feedback loops^5–11^. However, not all genes that physically interact with each other regulate each other. Interaction does not necessitate regulation. For example, the mRNAs bound by a given RBP and the RNAs that change expression upon deletion of that RBP show surprisingly little overlap^12^. Therefore, high-throughput experimental and computational methods that are increasingly good at correctly identifying physical interactions must be complemented by high-throughput methods for quantifying the sign and strength of these regulatory interactions.

Feedback loops can be described by two sets of properties: positive or negative and direct or indirect^13,14^. A direct auto-regulatory feedback loop is one in which a protein activates or inhibits itself, while in an indirect loop this feedback occurs through one or more intermediate genes. Furthermore, feedback loops can be positive, in which a gene increases its own expression or activity (eg: via phosphorylation), or negative, in which a gene represses or inactivates itself. Direct auto-regulatory negative feedback loops have an intrinsic ability to reduce sensitivity to intrinsic and extrinsic perturbations ^15–17^. In addition, the negative feedback network motif can shorten the response time of a network, as is found in the SOS DNA repair pathway and in ribosome biogenesis in E coli^18–20^, and can alter the response curve of a gene to changes in inducer concentration^21,22^. Time-separation in negative feedback loops, often in indirect feedback via intermediate genes, can generate oscillations and irreversible transitions, as are found in the circadian clock and the cell-cycle^23,24^. Like negative feedback, positive feedback can produce sustained oscillations^25^. However, biologically, positive feedback loops play an entirely different set of roles^26^, the most common of which is bi-stability, or all-or-none-transitions^27^. These networks motifs are common in the cell-cycle and in cell-differentiation, both of which often display a fast positive feedback loop and a delayed negative feedback loop in order to achieve an irreversible transition between two stable states, such as in the cell-cycle or in sex differentiation^24,28^.

Functional genomic and bioinformatic methods predict that many RNA binding proteins (RBPs) participate in feedback loops, in which the RPB regulates the level of its own protein, either directly or indirectly^10^. Direct negative feedback auto-regulation appears to be a common motif for RBPs, and many genes that lack a canonical RNA binding domain may bind to their own mRNA and regulate their own translation^10^. However, very few of these feedback interactions have been experimentally tested, as no high-throughput methods exist for such validation. The typical high test to measure the regulatory effect of a particular RBP involves deletion of that RBP, and therefore these methods are incapable of determining the existence of feedback loops. In order to determine the complete set of RBPs that control their own expression via direct or indirect feedback regulation, we developed a mathematical model and a high-throughput experimental method that work together to identify the presence of such loops and to measure their relative strength.

## RESULTS

### A mathematical model for detecting feedback loops using a synthetic inducible promoter

In a simple feedback-free model of gene expression (see methods), two proteins under the control of the same inducible promoter will show similar inductions curves **(Figure 1A-D)**. Differences in the transcription, translation, and degradation rates of these genes result in offset induction curves **(Figure 1B)**. However, the offset between these two curves is constant, and therefore the log-ratio of expression between a protein of interest and a reference protein (eg: GFP) under control of the same promoter will remain constant across induction levels (**Figure 1C)**. This effect can also be visualized by plotting expression of the protein of interest against the reference; a change in the transcription rate, translation rate, or degradation rate will result in diagonal lines with the same slope (**Figure 1D)**. However, if one of the proteins participates in a feedback loop, the shape of the induction curve will change (**Figure 1E,F)**. At low levels of induction (low TF concentration, and therefore low protein levels), the feedback loop will be negligible, and expression will be similar between the two proteins. As the induction increases, and therefore the expression of both proteins increases, the effect of the feedback loop on expression will increase, resulting in larger differences in the ratio between the expression of the two proteins **(Figure 1G)**. This can also be visualized as a change in the slope when comparing the expression of one protein against the other **(Figure 1H)**. Therefore, a relatively simple model of gene expression suggests that it should be possible to detect feedback loops by placing two proteins under the control of the same inducible reporter, and determining how the ratio between the two proteins changes as a function of expression level.

**Figure 1.**
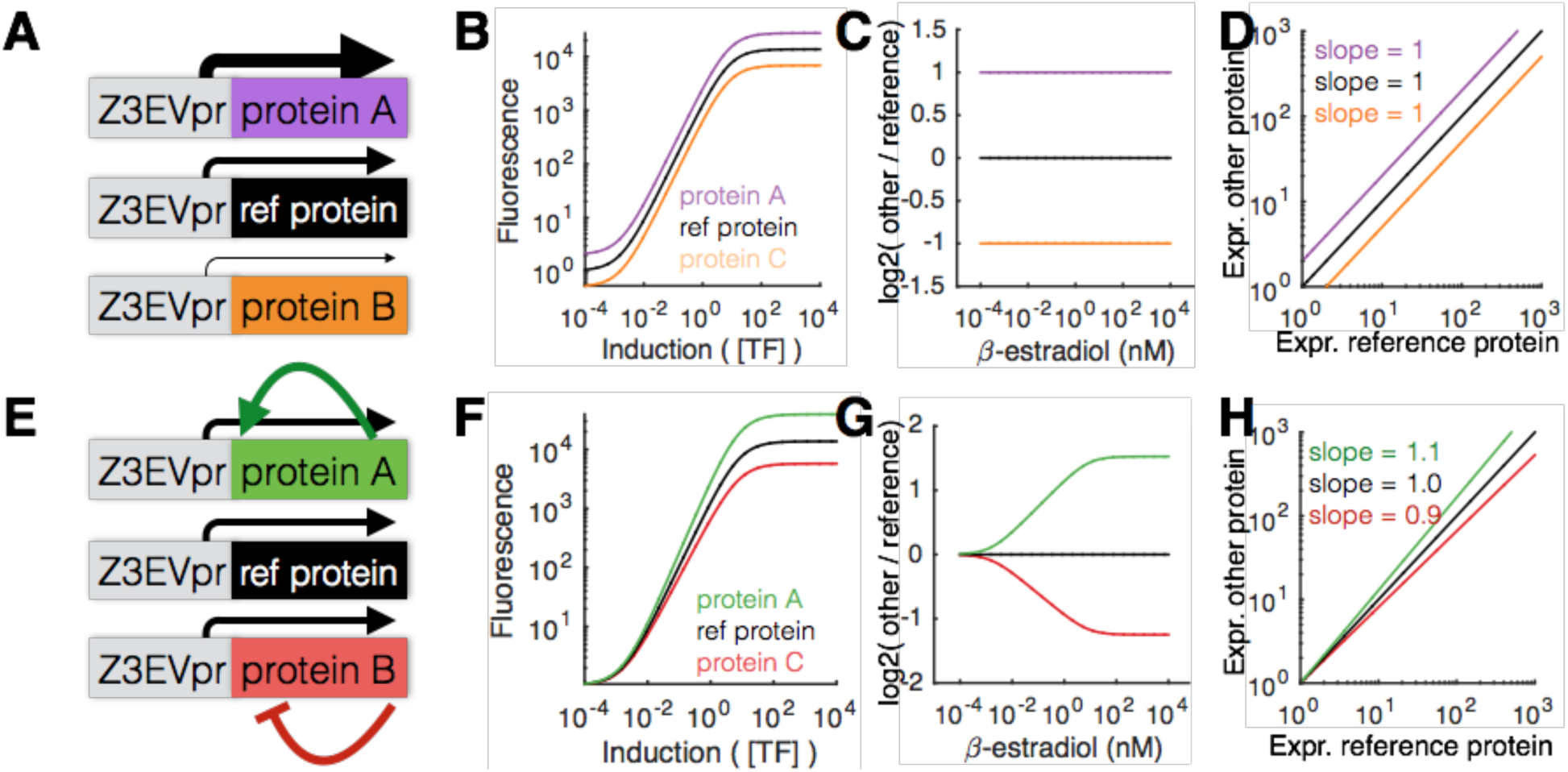
A mathematical model for detecting feedback loops using inducible promoters. **(A)** In a simple no-feedback model in which expression of three proteins is driven by identical promoters, but these genes differ in other characteristics, such as mRNA or protein stability, **(B)** all three proteins will be expressed with offset but identically shaped induction curves. **(C)** Therefore the log2 ratio of the expression level of a protein of interest and a constant reference protein will remain constant throughout the induction curve. **(D)** This can also be quantified by graphing the expression of each protein against a common reference; the lines will be offset from the diagonal but have the same slope. **(E)** In a model in which the expression of the genes of interest differs due to the presence of positive (green) or negative (red) feedback loops, **(F)** the induction curves will differ but not in a purely offset manner. **(G)** Instead, the log2 ratio of expression will change as a function of induction, because the effect of the feedback loop will be greater at greater expression levels. **(H)** If each protein of interest is graphed against a common reference, the result is that a positive feedback loop will result in a larger slope, while a negative feedback loop will result in a smaller slope.

### Design of a feedback-detector master strain

In order to experimentally test the above predictions, we developed a feedback detector control strain (*bud9*::Z_3_EVpr-GFP-GAL80_3’UTR,_ *his3*::Z_3_EVpr-mCherry-HIS3_3’UTR,_hereafter referred to as Y197) in which both GFP and mCherry are driven by the same Z_3_EV promoter under the control of the *β*-estradiol inducible synthetic transcription factor Z_3_EV **(Figure 2A,B)**. Addition of *β*-estradiol results in increased activity of the Z_3_EV transcription factor and increased expression of both GFP and mCherry (**Figure 2C)**. In the absence of feedback, the two proteins show identical induction curves, and the ratio between the two proteins remains constant (**Figure 2C,D**). Our model (see Methods) predicts that if we use the Z_3_EVpr-mCherry construct to N-terminally tag an RNA binding protein that participates in a feedback loop **(Figure 2E,F)**, we would alter the induction curve of mCherry but not GFP, and hence the ratio between the two fluors would change as a function of TF concentration (**Figure 2G,H)**. In order to experimentally test our feedback model, we built synthetic gene circuits with and without feedback regulation.

**Figure 2.**
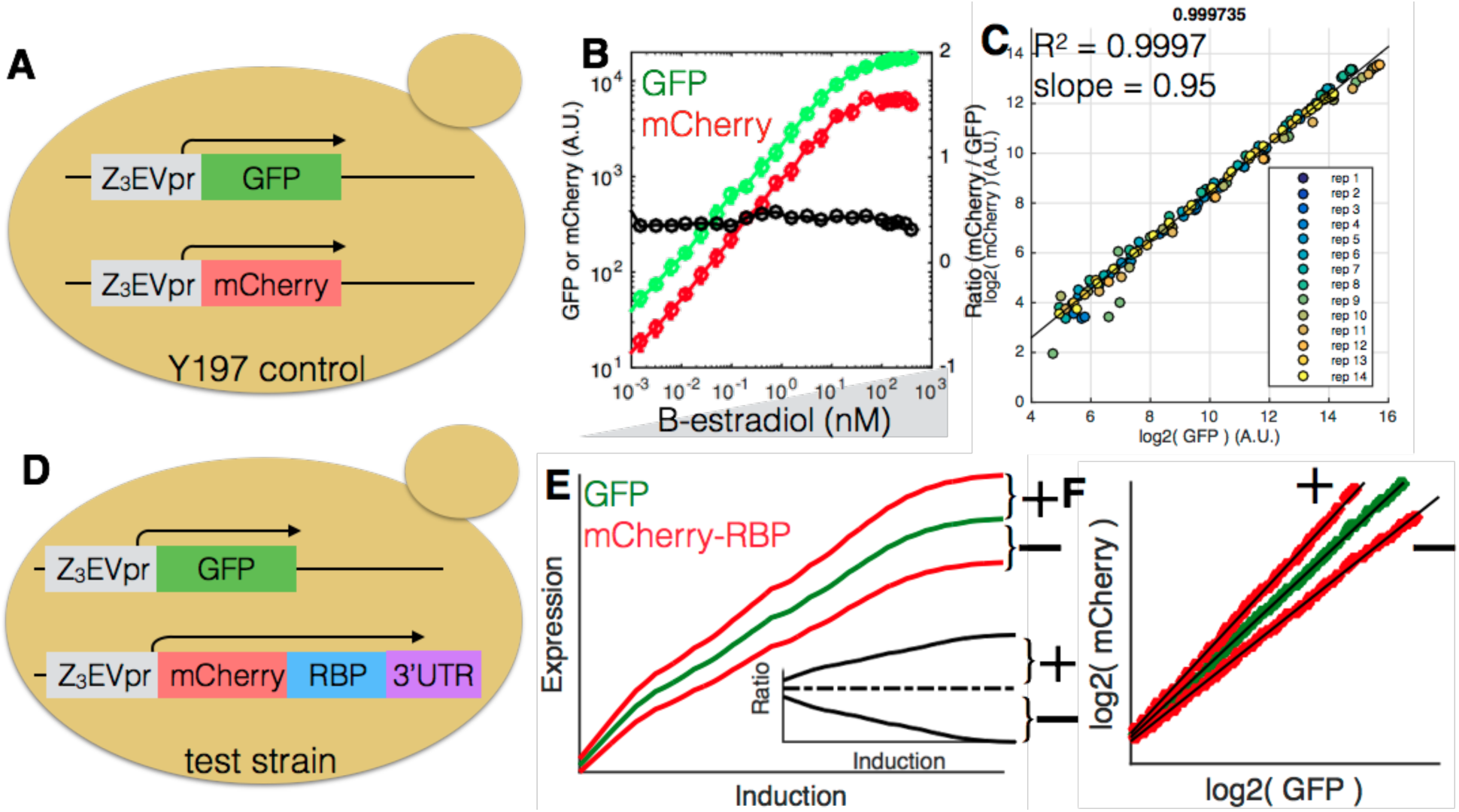
An experimental system to implement a feedback detector for RNA bindingproteins. **(A)** A control strain in which both GFP and mCherry are driven by the same β-estradiol inducible promoter, the Z_3_EVpr, which is a mutated GAL1 promoter that contains six binding sites for the synthetic zinc finger transcription factor Z_3_EV. **(B)** Both GFP and mCherry show similarly shaped offset induction curves, resulting in a relatively flat log2 ratio of expression between the two proteins. **(C)** Expression data from Y197 (panel A) are well fit by a straight line with a slope close to one, suggesting that GFP and mCherry show almost identical changes in expression across a wide range of β-estradiol concentrations. **(D)** A test strain for determining if an RBP participates in a feedback loop. The native RBP is replaced with an N-terminally tagged version driven by the Z3EV inducible promoter, in a strain that already contains Z3EVpr-GFP. Thus expression of the RBP can be monitored over a wide range of induction levels. **(E)** Simulated data using the measured GFP expression from Y197, showing the effect of positive or negative feedback on the induction curves and the log2 ratio between GFP and mCherry. **(F)** Given the measured GFP expression of Y197, the expected mCherry signal in the presence of positive or negative feedback.

### A Puf3-COX17 construct participates in a auto-regulatory negative feedback loop

Puf3 is an RNA binding protein that binds to sequence elements in the COX17 3’UTR, resulting in a destabilized COX17 mRNA^29^. We therefore hypothesized that a Z_3_EVpr-mCherry-Puf3-COX17_3’UTR_ strain (LBCY209, hereafter referred to as PUF3_COX17_) should have a negative feedback loop, while a Z_3_EVpr-mCherry-Puf3-PUF3_3’UTR_ (LBCY200, PUF3_PUF3_) should lack feedback regulation. We therefore built these two strains and measured the expression of mCherry-Puf3 as a function of *β*-estradiol. We find that, as predicted by the model, PUF3_PUF3_ shows the same induction curves as Y197, albeit shifted towards lower mCherry expression (**Figure 3B,C)**. In contrast, the mCherry signal for the PUF3_COX17_ flattens out at high *β*-estradiol (**Figure 3B,C)**. The result is that the the log2(mCherry/GFP) ratio shows identical behavior between Y197 and PUF3_COX17_ **(Figure 3D)**. In contrast, for PUF3_COX17_, the ratio decreases with increasing *β*-estradiol, consistent with the presence of a negative feedback loop. Further consistent with the presence of a negative feedback loop in PUF3_COX17_ but not in PUF3_PUF3_, the slope of GFP vs mCherry for the PUF3_COX17_ strain is lower **(Figure 3E)**, as predicted by the model for a negative feedback loop.

**Figure 3.**
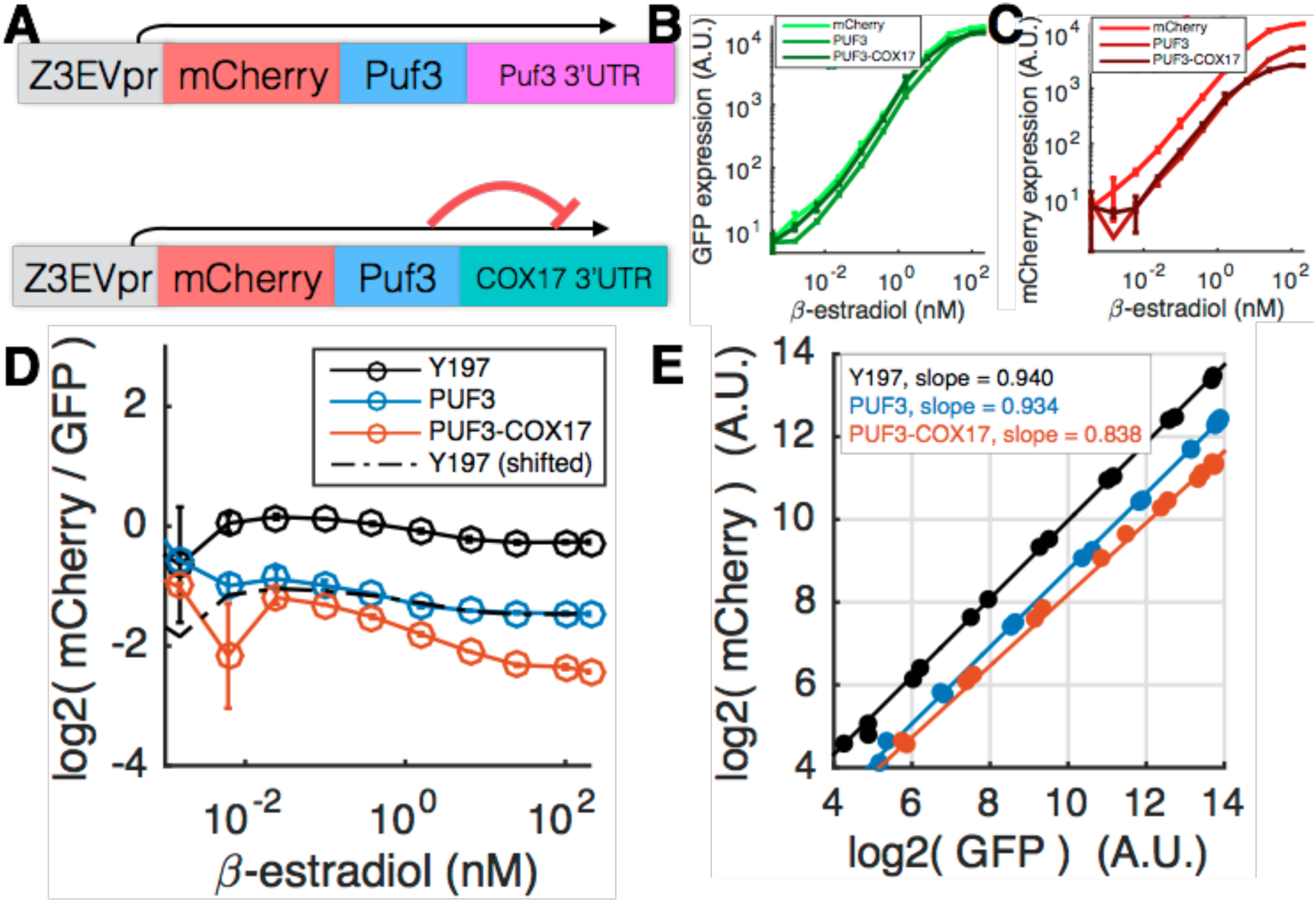
The RBP Puf3 with a COX17 3’UTR is a direct negative feedback loop. **(A)** A yeast strain in which the native Puf3 locus was replaced with either Z_3_EVpr-mCherry-PUF3 or Z3EVpr-mCherry-PUF3-COX17_3’UTR_. The latter strain is expected to generate a direct negative feedback loop, as Puf3 binding destabilizes the COX17 3’UTR. **(B)** GFP induction curves are similar in Y197, mCherry-PUF3 and mCherry-PUF3-COX17 strains. **(C)** The mCherry-Puf3 induction curve is offset from the Y197 mCherry induction curve, yet the two curves are similarly shaped. In contrast, the mCherry-Puf3-COX17 induction curve flattens out at high induction levels. **(D)** The log2 mCherry/GFP ratio shows a large decrease with increasing induction for PUF3-COX17, consistent with a negative feedback loop. mCherry-Puf3 alone has the same trend in ratio as does mCherry alone, consistent with no feedback loops. **(E)** The GFP vs mCherry slope of mCherry-Puf3 and mCherry are nearly identical. The Puf3-COX17 slope is lower, suggesting the presence of a negative feedback loop.

Gene expression is a stochastic process in which promoters switch on and off, generating bursts of protein production^30^. Therefore gene expression can be decomposed into two parts, burst frequency (the rate at which a promoter switches on), and burst size (the number of protein molecules made each time a promoter switches on)^31^. We hypothesized that a negative feedback loop that acts at the level of mRNA stability would decrease burst size. Consistent with this hypothesis, the burst frequency for the two strains is identical, while the burst sizes are vastly different **(Figure 4A-D)**. Interestingly, we find that, at low induction levels, PUF3_COX17_ has a higher burst size than PUF3_PUF3_, though the burst size of PUF3_COX17_ increases more slowly than that of PUF3_PUF3_**(Figure 4B)**, suggesting that two competing processes control expression of PUF3_COX17_: an increase in expression from increasing *β*-estradiol, and a decrease in expression from decreasing stability of the mRNA due to negative feedback. The final result of the negative feedback loop is that the same mean expression is reached at different *β*-estradiol concentrations, though the width of the single-cell expression distribution of mCherry-Puf3 is wider without the negative feedback loop**(Figure 4E)**, suggesting that cells may use auto-regulatory negative feedback loops to decrease cell-to-cell variability in RBP expression.

**Figure 4.**
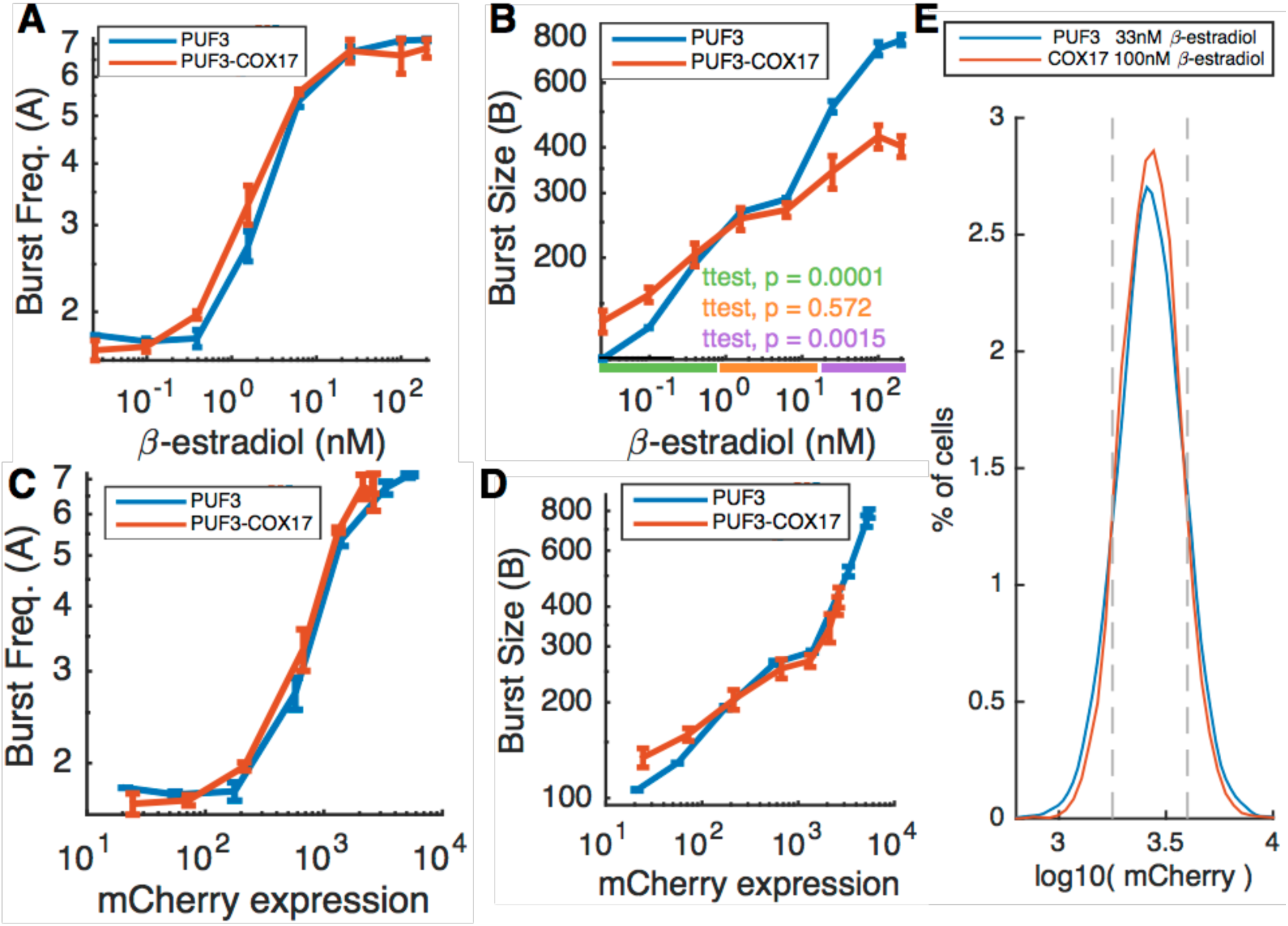
The PUF3FCOX17 negative feedback loop results in a change in burst size withincreasing Puf3FCOX17 expression. Burst frequency (A) and burst size (B) were inferred by fitting a gamma distribution to the fluorescence data. **(A)** Burst frequency increases similarly with increasing induction for both strains, suggesting that the negative feedback loop does not affect the frequency of transcriptional bursts. **(B)** Increasing β-estradiol results in increased burst size for both strains. Burst size increases more slowly for the PUF3-COX17 strain, suggesting that, as Puf3 expression increases, RNA stability, and therefore burst size, decreases. The difference in burst size is significantly different at both low and high induction levels. **(C)** The same data as in (A) but burst frequency graphed against expression. **(D)** The same data as in (B) but burst size graphed against expression. **(E)** There exist induction levels at which both strains have the same measured mean expression, but PUF3-COX17 shows a decrease in the width of the distribution.

### Z_3_EVpr-Pub1 participates in an indirect positive feedback loop

Pub1 is a poly(U) binding protein that binds to and stabilizes up to 10% of yeast mRNAs. We found that a *pub1Δ* strain exhibits reduced Z_3_EVpr-GFP expression **(Figure 5A)**. However, this reduction is not constant **(Figure 5B)**, as would be expected from altered stability of the GFP mRNA. Instead the effect depends on the level of induction, suggesting that Pub1 acts upstream of GFP, *i.e.*, at the level of Z_3_EV activity. This suggests that Pub1 may increase expression all Z_3_EV targets in a dose-dependent manner. In other words, a Z_3_EVpr-Pub1 will generate a positive feedback loop. To test this hypothesis we generated a Z_3_EVpr-mCherry-Pub1 strain **(Figure 5C)** and measured both GFP and mCherry as a function of *β*-estradiol concentration. Consistent with our hypothesis, both GFP and mCherry show more steep induction curves in a Z_3_EVpr-mCherry-Pub1 strain than they do in the wild-type control **(Figure 5D,E,F)**. In contrast to the negative feedback loop mediated by Puf3, the positive feedback loop mediated by Pub1 results in an increase in the width of the single-cell distribution **(Figure 5G)**. Thus, Z_3_EVpr-mCherry-Pub1 drives a positive feedback loop that results in increased expression of both Z_3_EVpr-mCherry and Z_3_EVpr-GFP.

**Figure 5.**
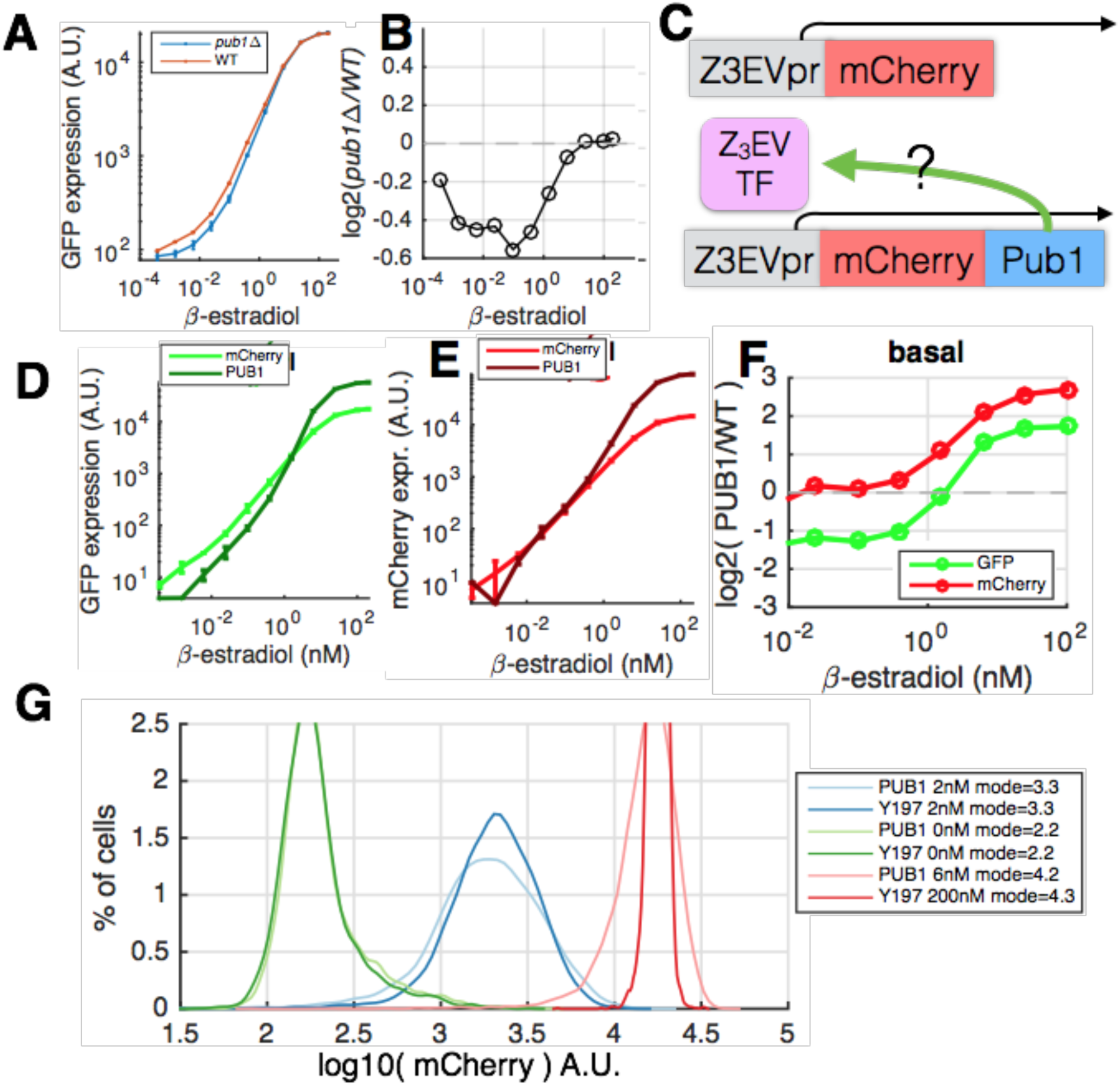
Z_3_EVprFPub1 generates an indirect positive feedback loop that acts on the Z_3_EVtranscription factor. **(A)** A *pub1Δ* strain shows decreased Z3EVpr-GFP expression at low β-estradiol concentrations. **(B)** The log2 ratio of GFP expression in *pub1Δ* vs wild-type cells shows that both strains have the same maximal GFP expression. **(C)** An inducible Pub1 strain for determining if this is due to regulation of the Z_3_EV TF by Pub1. **(D)** Z_3_EVpr-GFP exhibits a steeper induction curve in a Z_3_EVpr-mCherry-Pub1 strain. **(E)** Z_3_EVpr-mCherry-Pub1 exhibits a steeper induction curve than does Z_3_EVpr-mCherry in the Y197 control strain. **(F)** The log2(Z_3_EVpr-Pub1/WT) ratios for both GFP and mCherry increase with increasing β-estradiol, consistent with mathematical models of a positive feedback loop. **(G)** A positive feedback loop results in increased width of the single-cell expression distribution in the Z_3_EVpr-Pub1 strain.

### A mathematical model in which tagging RBPs with mCherry can explain all of the data

In the above experiments we measured expression of four different strains, all of which contain Z_3_EVpr-GFP as an internal control, and each of which contains a unique Z_3_EVpr-mCherry derived construct. The different C-terminal ends on mCherry (mCherry, mCherry-Puf3 and mCherry-Pub1), and the different 3’UTRs are likely to affect the stability of the mCherry mRNA and protein. These can be modeled as constant, transcription factor independent changes in mCherry expression level. In contrast, feedback loops introduced transcription factor dependent changes in mCherry expression. To determine if all of our data can be explained by this model, we fit a model without feedback to Y197 (mCherry) data, and then used the same parameters, but allowed only K’_b_ (TF independent transcription & translation) to vary (see methods). We find that this model can fit Y200 (mCherry-PUF3_PUF3_) but not the other strains, consistent with our above hypothesis that this strain lacks any feedback loops **(Figure 6A,B)**. We next fit the same model but allowed both K’_b_ and F to vary. We find that this model can fit all measured strains, and that all good fits to Y209 (mCherry-PUF3_COX17_) have a negative value of F, and all good fits to (mCherry-PUB1) have a positive value of F, consistent with our hypothesis that these strains have negative and positive feedback loops, respectively **(Figure 6C,D)**. Thus, our two-color Z_3_EVpr feedback detector system is able to identify the presence of both direct and indirect feedback loops regulated by RNA binding proteins.

**Figure 6.**
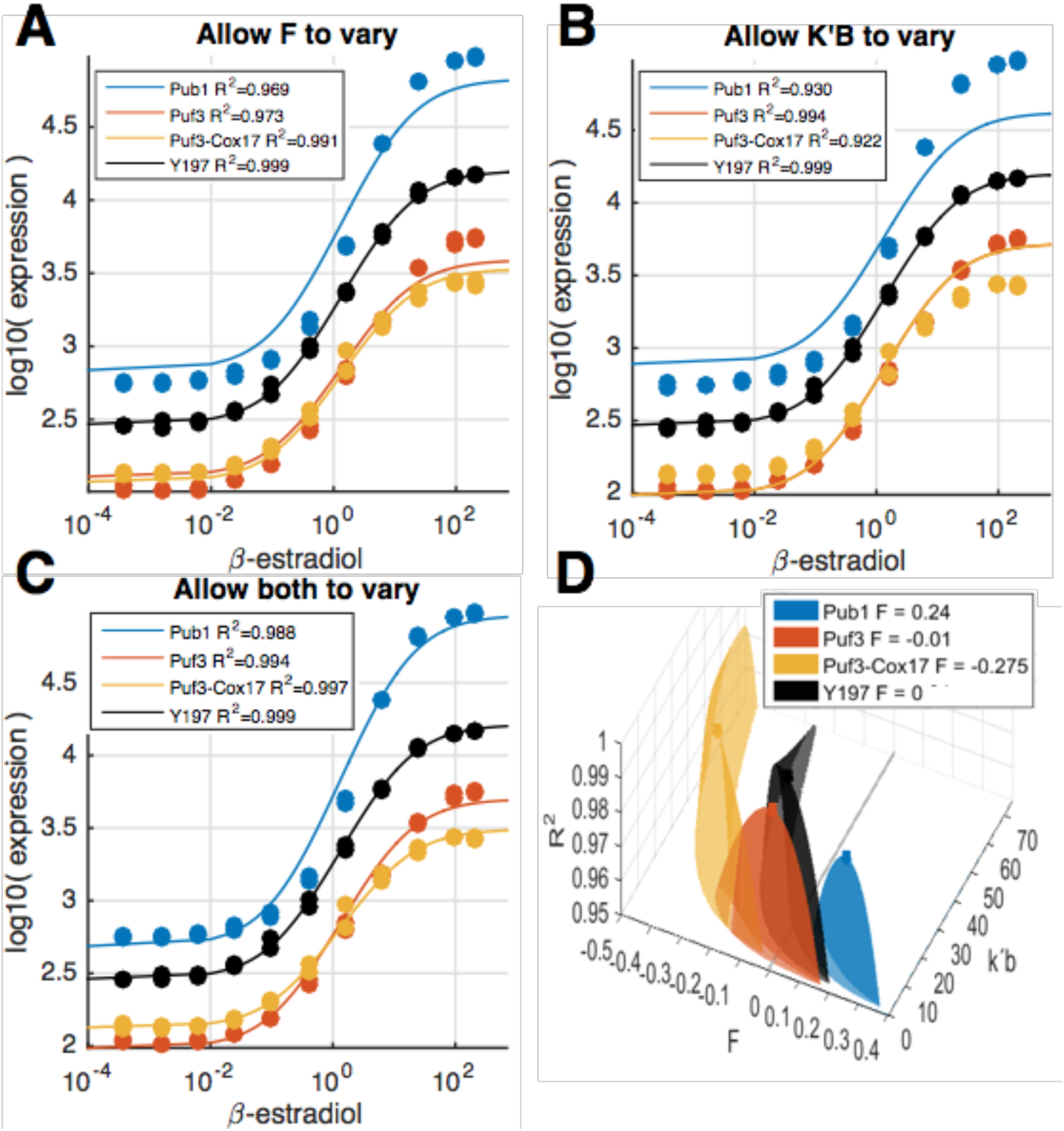
A mathematical model including feedback is required to fit the Z_3_EVpr-Pub1 and Z_3_EVpr-Puf3-COX17 data, but not the Z3EVpr-Puf3-PUF3 data. A model that does not incorporate feedback was fit to mCherry expression data from Y197, and either one or two parameters were allowed to vary to fit data from the other strains. **(A)** A model fit to Y197 in which only the feedback parameter F is allowed to vary can fit fit all strains with an R^2^ greater than 0.95. **(B)**A model in which only k’b is allowed to vary can only fit PUF3 with an R^2^ greater than 0.95. **(C)** A model in which both F and k’b are allowed to vary is able to fit all experimental strains. **(D)** The effect of varying both F and k’b on the ability of a model fit on Y197 to fit the other strains is shown. Only R^2^ values >= 0.95 are shown for clarity.

## DISCUSSION

In summary, we have developed and experimental and mathematical framework to measure the strength of native regulatory feedback loops. Through the use of an experimental system in which two fluorescent reporters are driven by the same transcription factor, but one of these is coupled to an RNA binding protein, we can not only detect the presence of both positive and negative feedback loops, but we can quantitatively measure the strength of these loops as well. We applied this system to a synthetic auto-regulatory negative feedback loop with the RBP Puf3, and found that introduction of this loop reduces the noise in Puf3 expression. In addition, we applied this system to the RBP Pub1, and detected a positive feedback loop that increases expression noise. However, this feedback loop is not specific to Pub1, but acts on both Z_3_EVpr-mCherry-Pub1 and Z_3_EVpr-GFP, suggesting that it acts upstream of Pub1, possibly by directly regulating the concentration of the Z_3_EV TF. Pub1 binds the ENO2 3’UTR and the ACT1 5’UTR^12,32^. This observation highlights an important strength of our dual-reporter method, as opposed to more traditional approaches in which the gene of interest is overexpressed or deleted without an internal control in the same cell. Many genetic perturbations result in both direct and indirect effects. Varying Pub1 changes both mCherry and GFP, showing that the effect is non-specific. The vast majority of changes in mRNA levels observed in deletion and overexpression experiments are indirect^33^. GFP serves as an internal control for the state of the cell and for the state of the synthetic gene circuit. Thus, by combining the dual reporter system with a mathematical model we can accurately different direct and specific effects from indirect pathway-specific or global effects.

Consistent with past theory and experiments^15,17,34–37^, we find that a negative feedback loop decreases noise, while a positive feedback loop increases noise. Interestingly, the increase in noise is far larger than the decrease in noise. However, at this point we cannot say if this difference in the magnitude of the changes is a general property of positive vs negative loops or of direct auto-regulatory loops vs pathway-specific or global feedback loops. It will be interesting to determine if global regulators of expression, such as Pub1, also act as global regulators of expression noise, or if the large increase in noise is due to a pathway-specific positive feedback loop.

Interestingly, we find that the parameter regime that fits Z_3_EVpr-mCherry-Puf3 is exactly continuous with the regime that fits Z_3_EVpr-mCherry-Puf3-COX17 **(Figure 6D)**. In addition, the induction curves intersect at around the position of half-maximal expression **(Figure 6C)**. This suggests that, at half-maximal induction, the two constructs have identical transcription and translation rates, but that at low induction the COX17_3’UTR_ mRNA is more stable (less Puf3), while at high induction the reverse is true.

Finally, this system may serve as a platform for the design and characterization of synthetic RBPs^38,39^. Synthetic regulatory circuits with designed sequence specificities have many advantages over the repurposing of native circuits, such as Gal4-UAS^40^. However, it is difficult to design such circuits so that they do not interfere with the host cell^41^. The dual-reporter aspect of our system ensures that secondary effects of synthetic circuits can easily be detected, and constructs that lack secondary effects chosen.

## MATERIALS AND METHODS

### Yeast strains and media

All yeast strains are listed in **Supplementary Table 1**. As non inducible and autofluorescence control we used FY4, a wild prototrophic yeast strain^42^. The parental strain for all Z3EV strains is DBY19054^43^. To generate LBCY197, we generated a PCR amplicon containing KanMX-Z3EVpr-mCherry using primers 196 & 197 and LBCP80, and transformed this amplicon into DBY19054. To generate yeast strains LBCY200 and LBCY201 we first generated plasmids LBCP94 and LBCP95, and amplified these plasmids using primer pairs 325,326 and 365,366 respectively, and transformed these PCR amplicons into DBY19054. In order to generate LBCY209 we first generated a puf3::HYG^R^ strain (LBCY203), and then generated LBCY209 using primers 325 & 327 and plasmid LBCP96. Colony PCR was used to confirm correct integrations of all strains, and to verify that the Z_3_EV promoter has the the correct number of Z_3_EV binding sites. All transformations were performed using the standard lithium acetate method^44^. PCR for transformation was performed with Phusion DNA Polymerase (*Sigma Aldrich*). Colony PCR was performed using Taq Polymerase 2x Master Mix (*Sigma Aldrich*). Selection for drug resistant transformants was done on YPD plates with Hygromycin B(IBIAN), CloNAT(Werner bioreagents), or G418(VWR).

### Plasmid construction

In order to create plasmid LBCP80, a 1.5kb PCR amplicon containing the *Z_3_EVpr* from gDNA of DBY19054 was amplified using two rounds of PCR, first with primers 316 & 317, then with primers 308 & 309. This PCR product, along with mCherry, was cloned by Gibson assembly into plasmid PYM-N14^45^ which had cut with SacI and EcoRI to remove the GPD promoter. To create plasmids LBCP94 & LBCP95 the PUF3 and PUB1 ORFS were PCR amplified using primer pairs 318, 319 and 329, 330 and Gibson cloned into EcoRI linearized LBCP80. Plasmid LBCP96 was created using Gibson assembly with EcoRI linearized LBCP80, the PUF3 ORF, and a 200 bp 3’UTR region of COX17 PCR amplified from the genome using primers 321 & 322. After Gibson assembly, each plasmid was transformed to *E. coli* by electroporation and transformants confirmed by colony PCR, minipreped and checked by Sanger-sequencing and multi-site restriction digest.

### Flow Cytometry

Single colonies were picked from YPD plates and cultured overnight in SCD media, inoculated at OD_600_ = 0.02 into different concentrations of SCD + *β*-estradiol (Sigma E8875) and measured after 7.5 h of growth at 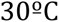 (in which OD_600_ was between 0.25 and 0.5). The flow cytometry machine used was BD LSRFortessa (BD Biosciences) and with 488nm and 561nm lasers with 530/28 or 610/20 filters for GFP and mCherry. All data analysis was performed using MATLAB as previously described^4^.

### Mathematical modeling and fitting to data

To construct a mathematical model that could explain the fluorescence values over the range of inducer concentrations, it was assumed a constant rate of mRNA synthesis and a mRNA degradation rate proportional to the actual mRNA concentration, and a protein synthesis rate proportional to the mRNA concentration and a degradation rate proportional to the actual protein concentration. Thus, it was defined the following ODE system, which defines the background model

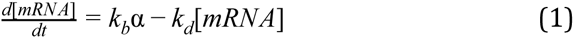

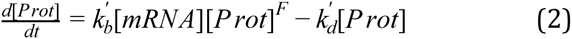

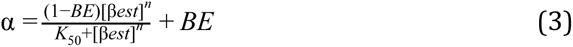

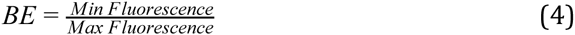

in which k_b_ and k’_b_ are rates of transcription and translation, respectively, and k_d_ and k’_d_ rates of mRNA degradation and protein degradation, respectively. The factor ɑ is the transfer function that establishes the relation between inducer concentration and promoter activation, which has been supposed to follow a Hill equation. K_50_ is the concentration of *β*-estradiol at which expression reaches its half-maximal value. F is a feedback constant, with negative values for negative feedback interactions and positive values for positive feedback. Considering (1) and (2) at equilibrium, we can write

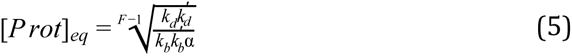

We note that this feedback model is invalid with regards to the rate of protein production when [Prot] approaches 0, which is not what we would expect for such a biological system. We therefore used fluorescence data, which always has values greater than zero, as a proxy for [Prot]. We note that all tested strains show GFP and mCherry signals significantly above background in the absence of *β*-estradiol, suggesting that, averaged across the population, [Prot] is always greater than zero. Furthermore, we note that each of the variables within (k_d_ *k’_d_)/(k_b_ *k’b), cannot measured individually using our system. We therefore vary a single of these parameters and keep the other three constant.

All data were first normalized by subtracting either autofluorescence, as measured from a strain lacking GFP and mCherry, or basal expression, as measured at 0nM B-estradiol. All analysis, figures, and model fitting was performed using both normalization methods; the results are qualitatively identical. The latter corrects for differences in the basal expression level between strains, and is used when plotting GFP vs mCherry. The former explicitly shows differences in basal expression level and is used for fitting the model to data. Prior to fitting, normalized fluorescence measurements were log10 transformed to prevent the high expression values from dominating the fit. The model was first fit to data from Y197. Then, either F, k’b, or both F and k’b were varied over two orders of magnitude and the R^2^ was calculated between model and data.

In order to quantitatively decide what R^2^ constitutes a good fit, for each strain, we fit the model to one biological replicate and then calculated R^2^ of that model a different biological replicate of the same strain. The R^2^ is always greater than 0.95 for all strains **(Supplementary Table 2)**; we therefore chose 0.95 as the threshold.

## ACKNOWLEDGEMENTS

We’d like to thank Gian Tartaglia, Marçal Gabaldá and Elena Abad for useful discussions. This work was supported by startup funds from the department (DCEXS) and grant from the Agència de Gestioó d’Ajuts Universitaris i de Recerca (AGAUR) to L.B.C. M.A.S.T was supported by a Evolutionarly Biology and Complex Systems (BESC) Program undergraduate fellowship. The funders had no role in study design, data collection and interpretation, or the decision to submit the work for publication.

## AUTHOR CONTRIBUTIONS

The author have made the following declarations about their contributions: Conceived and designed the experiments and wrote the paper: LBC, with help from MAST & JDE. Performed the experiments and analyzed the data: MAST, CTO, LBC. Contributed reagents and materials: MAST, CTO, JDE, LE.

